# Three-dimensional multilayer concentric bipolar electrodes enhance the selectivity of optic nerve stimulation

**DOI:** 10.1101/2022.03.21.485100

**Authors:** Eleonora Borda, Vivien Gaillet, Marta Jole Ildelfonsa Airaghi Leccardi, Elodie Geneviève Zollinger, Ricardo Camilo Moreira, Diego Ghezzi

**Affiliations:** Medtronic Chair in Neuroengineering, Center for Neuroprosthetics and Institute of Bioengineering, School of Engineering, École Polytechnique Fédérale de Lausanne, Switzerland

## Abstract

**Objective:** Intraneural nerve interfaces often operate in a monopolar configuration with a common and distant ground electrode. This configuration leads to a wide spreading of the electric field. Therefore, this approach is suboptimal for intraneural nerve interfaces when selective stimulation is required.

**Approach:** We designed a multilayer electrode array embedding three-dimensional concentric bipolar electrodes. First, we validated the higher stimulation selectivity of this new electrode array compared to classical monopolar stimulation using simulations. Next, we compared them in-vivo by intraneural stimulation of the rabbit optic nerve and recording evoked potentials in the primary visual cortex.

**Main results:** Simulations showed that three-dimensional concentric bipolar electrodes provide a high localisation of the electric field in the tissue so that electrodes are electrically independent even for high electrode density. Experiments in-vivo highlighted that this configuration leads to evoked responses with lower amplitude and more localised cortical patterns due to the fewer fibres activated by the electric stimulus in the nerve.

**Significance:** Highly focused electric stimulation is crucial to achieving high selectivity in fibre activation. The multilayer array embedding three-dimensional concentric bipolar electrodes improves selectivity in optic nerve stimulation. This approach is suitable for other neural applications, including bioelectronic medicine.

## 1. INTRODUCTION

The treatment for several neurological disorders and traumatic injuries, such as Parkinson’s disease, epilepsy, deafness, blindness, amputation, chronic pain and spinal cord injury, among others, relies on the electrical stimulation of the central and peripheral nervous system [1]. Interfaces to the nervous system traditionally consist of microelectrode arrays placed on (epineural) or inside (intraneural) the tissue. Microelectrodes deliver current or voltage pulses to stimulate neuronal cells and recover lost functions.

The number of electrodes in the array varies depending on the application, from few units (e.g. deep brain stimulation) [2] to few tens (e.g. cochlear implants) [3], or even hundreds or thousands (e.g. retinal prostheses) [4,5]. Regardless of the number of electrodes, neural interfaces share the same principle: electrodes should be electrically independent without stimulation cross-talk among neighbours. In other words, each electrode should excite selectively a specific subset of cells or fibres, different from those stimulated by neighbouring electrodes. For example, a visual prosthesis should have as many electrically independent electrodes as possible so that each electrode induces a spatially isolated visual percept in the subject, called phosphene. Artificial vision consists of perceiving the world using an ensemble of these phosphenes [6]. On the other hand, a second requirement is to have electrodes as dense as possible to increase the theoretical resolution of the restored image (as the number of pixels per unit of area). Although these two concepts work in theory, there is a constraint in practice: a trade-off between density and independence must be found. Increasing the electrode density increases the risk of having a stimulation cross-talk among neighbouring electrodes. In such a case, two electrodes could induce the same phosphene, or two close phosphenes could fuse [7,8].

The configuration used for electrical stimulation plays a significant role in this balance. Most clinical neural interfaces function in a monopolar configuration with a common and distant return ground placed several millimetres away [9–12]. When the current is injected into the stimulating electrode, the electric field spreads to the common and distant ground, not allowing the confinement of the electrical stimulation around the electrode and resulting in the activation of a large portion of the neural tissue [13]. Other stimulation configurations were widely investigated in cochlear implants and visual prostheses, including bipolar, hexapolar and multipolar [14]. In addition to these configurations, the concentric bipolar (CB) is an attractive strategy. Each stimulating electrode has a concentric return electrode, and the electric field is localised between the two electrodes [14].

Compared to the monopolar configuration, the main disadvantage of a conventional CB configuration is that two feedlines are required for each electrode pair, composed of the stimulating electrode and the concentric return electrode. Two feedlines increase the space occupied on the array and, consequently, the size of the implant. This paper reports the design, simulation and validation of a novel intraneural electrode array based on three-dimensional (3D) multilayer CB electrodes fabricated using a previously described 3D multilayer technology [15]. This multilayer process allows placing electrodes on separate layers and overlaying the feedlines. Therefore, each electrode pair occupies the same space on the array as the monopolar configuration. This advantage is beneficial for intraneural interfaces where the nerve size is an important constraint to the total size of the array. Hence, we validated this technology in the context of intraneural optic nerve stimulation. Our research group recently proposed a new intraneural electrode array (OpticSELINE) for artificial vision via optic nerve electrical stimulation [16,17]. However, the selective activation of a few optic nerve fibres by each electrode was still an open challenge. The array of 3D multilayer CB electrodes proposed here improves the selectivity of the intraneural stimulation.

Beyond optic nerve stimulation and artificial vision, we believe that the high-spatial stimulation selectivity of this electrode array can be exploited in other neural applications, including bioelectronic medicine.

## 2. METHODS

### 2.1 Electrode array modelling

A finite element analysis (FEA) was performed in COMSOL Multiphysics (Version 5.3), using a stationary current study with the AC/DC module. The electrode array (5 × 5 electrodes, 80-μm diameter and 100-μm pitch) was simulated inside a 1-cm height and 2-cm wide cylinder filled with saline (saline box). A circular ground electrode (1-cm diameter) was placed concentrically with the simulated array (**Figure 1a**). The electrodes were modelled in platinum (Pt), while the substrate and superstrate were modelled in polyimide (PI). Meshing was performed with tetrahedral nodes. Simulations were performed under static conditions. Current conservation was assumed following the continuity equation. Electrodes were modelled with the geometrical and electrical parameters reported in **Table 1**.

**Figure 1.**
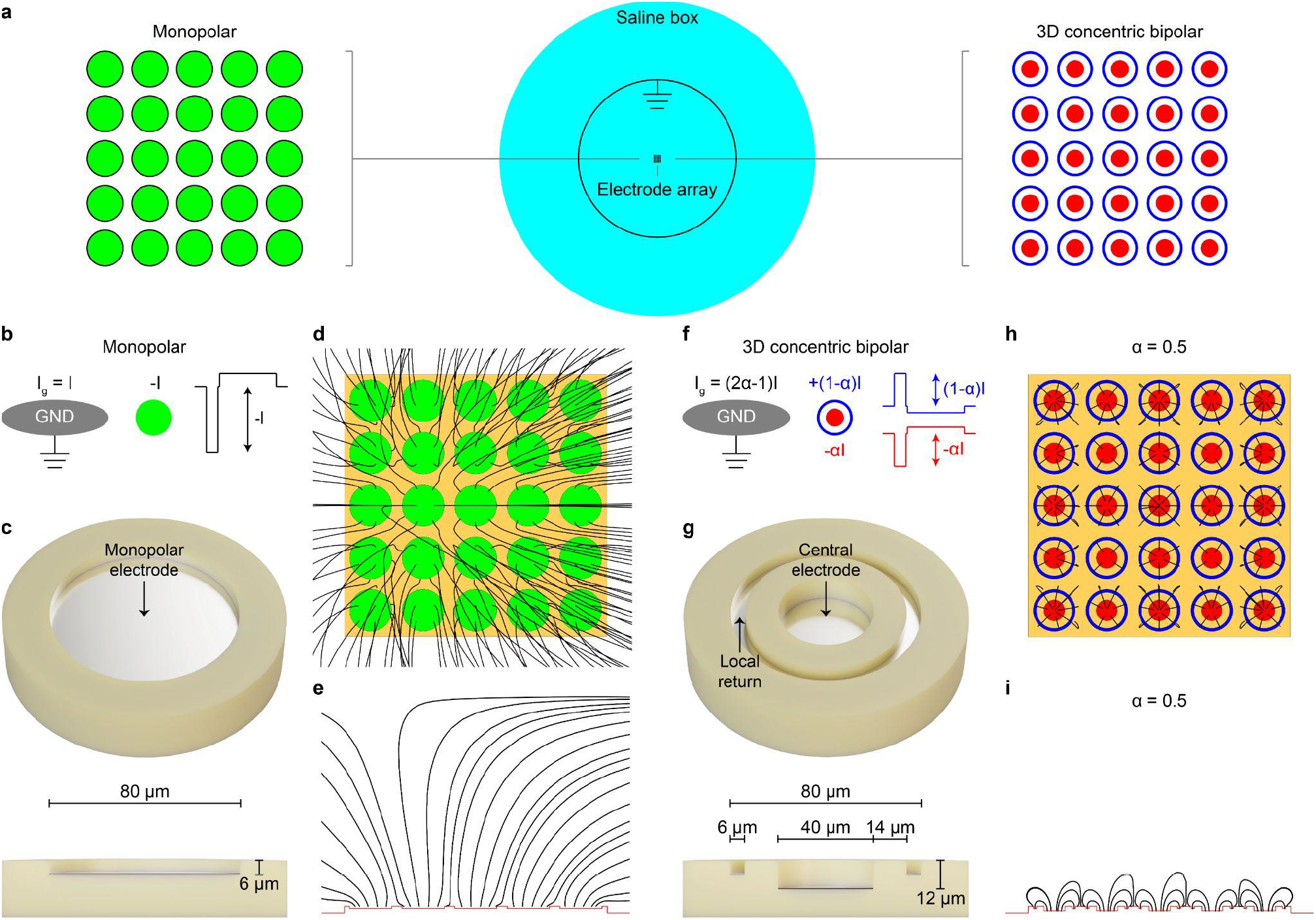
(**a**) Sketch of the FEA simulation. The electrode array (5 × 5 electrodes, 80-μm diameter and 100-μm pitch) is inside a 1-cm height and 2-cm wide saline box. A circular ground electrode (1-cm diameter) is placed concentrically with the electrode array. (**b**) Monopolar configuration: the current is injected into one electrode (negative for cathodic stimulation) with a faraway common ground. (**c**) Tilted top view (top) and side view (bottom) of a monopolar electrode. The encapsulation material (PI) is yellow, and the electrode (Pt) is grey. (**d**,**e**) Top view (**d**) and side view (**e**) of the electric field computed by FEA simulation for the monopolar configuration (all electrodes are active). (**e**) 3D multilayer CB configuration: the current is injected between the central electrode (cathodic, -αI) and the concentric return electrode (anodic, +(1-α)I), both referenced to the distal ground (I_g_ = (2α-1)I). (**g**) Tilted top view (top) and side view (bottom) of a 3D multilayer CB electrode. (**h,i**) Top view (**h**) and side view (**i**) of the electric field computed by FEA simulation for the 3D multilayer CB configuration (α = 0.5, all electrodes are active).

**Table 1.** Geometrical and electrical parameters for the model.

| Geometrical parameters |  |  |  |  |  |  |  |
| --- | --- | --- | --- | --- | --- | --- | --- |
|  | Central electrode |  |  | Return Electrode |  |  | Wall |
| | Diameter [$\mu\text{m}$] | Depth [$\mu\text{m}$] | Area [$\mu\text{m}^2$] | Width [$\mu\text{m}$] | Depth [$\mu\text{m}$] | Area [$\mu\text{m}^2$] | Width [$\mu\text{m}$] |
| **Monopolar** | 80 | 6 | 5027 | - | - | - | - |
| **3D multilayer CB** | 40 | 12 | 1257 | 6 | 6 | 1395 | 14 |
| Electrical parameters |  |  |  |  |  |  |  |
|  | Platinum |  | Polyimide |  | Saline |  |  |
| **Conductivity [$\text{S m}^{-1}$]** | $8.9 \times 10^6$ | | $6.67 \times 10^{-18}$ | | 1 | | |
| **Relative permittivity** | - |  | 3.4 |  | 80 |  |  |

### 2.2 Hybrid FEA-NEURON modelling of optic nerve stimulation

The hybrid FEA-NEURON simulation was performed as previously described [16]. The optic nerve model was built in COMSOL with the geometrical and electrical parameters reported in **Table 2**. The optic nerve and the meningeal layers were modelled as concentric cylinders. The axonal compartment was 1.5 mm in diameter. The pia was approximated by a contact impedance due to its small thickness, and cerebrospinal fluid was placed between the pia and the dura. A saline box (4.5-mm diameter and 5-mm length) surrounded the optic nerve. Except for the axons, which have higher longitudinal conductivity, the other domains have isotropic conductivity. A single monopolar or 3D multilayer CB electrode was modelled as described above and placed in the optic nerve centre. A ground electrode was placed in the saline box. The size of the saline box and the mesh were optimised to respect the hypothesis of ground condition at infinity. The mesh size was reduced close to the electrode to detect the rapid change in the electric field.

**Table 2.** Geometrical and electrical parameters for the optic nerve model. Saline conductivity was adjusted to 37 °C.

| Axons |  |  | Pia |  | Cerebrospinal fluid |  | Dura |  | Saline |
| --- | --- | --- | --- | --- | --- | --- | --- | --- | --- |
| Radius [$\mu\text{m}$] | Transversal conductivity [$\text{S m}^{-1}$] | Longitudinal conductivity [$\text{S m}^{-1}$] | Thickness [$\mu\text{m}$] | Conductivity [$\text{S m}^{-1}$] | Thickness [$\mu\text{m}$] | Conductivity [$\text{S m}^{-1}$] | Thickness [$\mu\text{m}$] | Conductivity [$\text{S m}^{-1}$] | Conductivity [$\text{S m}^{-1}$] |
| 750 | 0.08 | 0.5 | 10 | 0.016 | 100 | 1.7 | 300 | 0.06 | 1.7 |

The axon fibre model was implemented in NEURON (Version 7.4) as McNeal’s cable model. Briefly, only the nodes of Ranvier were active segments, while the myelinated segments were approximated by a perfect insulator. The geometrical parameters of the axon model were obtained from previous work [18]. Each axon model was a modified Hodgkin-Huxley model [19]. Each node of Ranvier contained five currents: fast sodium current, fast potassium current, persistent sodium current, slow potassium current, and leak current. The electrical properties of the axon model were obtained from previous work [20]. An axon is activated if an action potential travels to both ends. For an axon of a given diameter and shift (the relative distance separating its central node of Ranvier from the centre of the stimulating electrode), its probability of being activated by the stimulation was either 0 or 1. To determine the probability for a specific pair of coordinates in the nerve cross-section to contain an axon activated by a given electrical stimulus, 42 combinations of 6 shifts and 7 diameters were considered. Shifts ranged from 0, if the electrode was aligned with the central node of Ranvier, to 0.5, if the electrode was aligned with the centre of the myelinated segment, with steps of 0.1 since any shift is as likely as the others. On the contrary, the 7 diameters do not have the same probability of occurring: some will contribute more than others to the final activation probability. Instead of computing the activation probability for specific diameters with a constant interval and weighting these values by their frequency of occurrence, we computed the frequency distribution of axon diameters and multiplied it by the squared diameters. Then, we divided the distribution into 8 bins of 12.5% each, effectively doing the weighting before computing the probability values. Last, the 42 probability values were averaged to obtain the final activation probability by a given electrical stimulus for a given pair of coordinates in the nerve cross-section. For each current amplitude (2, 3, 5, 10, 20, 25, 50, 75, 100, 150, 200, 250, 500, 750, 1000, 1500 and 2000 μA) and configuration (monopolar and 3D multilayer CB) tested, an activation probability matrix was obtained by sampling every 40 μm × 40 μm pair of coordinates over the cross-section of the nerve. Then, they were sampled with higher resolution at the locations where a large change in the activation probability was present. The total number of fibres activated for each current amplitude and configuration was computed by multiplying each activation probability in the matrix by the number of fibres present in the sampled area (averaged fibre density in the rabbit optic nerve is 0.19 fibres μm^-2^, [16]) and summing up all the values. Activation probability maps were obtained by interpolating the activation probability matrix with a resolution of 1 x 1 μm^2^ using the cubic 2D-interpolation function in MATLAB (MathWorks).

### 2.3 Array microfabrication

The Flat-OpticSELINE [17] is a modified version of the OpticSELINE [16] without protruding wings. Therefore, it is equivalent to a transverse intrafascicular multichannel electrode (TIME) array. The Flat-OpticSELINE electrode array contains 8 CB electrode pairs fabricated with a 3D multilayer process. The detailed process flow is available in [15]. Briefly, a Ti/Al release layer (10/100 nm) was deposited using a magnetron sputter onto 4-inch Si wafers. The deposition of a PI layer (PI2611 HD MicroSystems GmbH) of 12 μm was obtained by spin-coating at 1000 rpm, soft-baking at 65 °C (5 min) and 95°C (5 min), as well as hard-baking at 200°C (1 hr) and 300°C (1 hr) both under nitrogen atmosphere. The electrodes Ti/Pt (15/300 nm) were made by magnetron sputtering onto oxygen plasma-treated PI layers, followed by photolithography and chlorine-based dry etching. The central electrodes were 40 μm in diameters with 15-μm wide interconnects. After photoresist removal, the substrates were coated again with 6 μm of PI (2000 rpm), as just described. The metallisation and PI encapsulation were repeated for the top layer, with a PI thickness of 6 μm. A 14-μm thick layer of positive photoresist was spin-coated and patterned by photolithography to open the electrodes and the pads for the connector in both layers. The PI layers were directionally dry-etched using oxygen plasma until the openings reached all the electrode layers (Pt was used as etch stop material). The remaining layer of photoresist was dissolved in acetone. The top layer electrodes had a 6-μm width ring shape, and 15-μm wide interconnects overlaid to the bottom ones. The Flat-OpticSELINE was then shaped by laser cutting and released from the wafer by Al anodic dissolution. After release, the arrays were inserted into a ZIF connector placed on a customised printed circuit board. Last, the electrodes were electroplated with platinum black (PtB), using a solution containing 1% of platinum chloride H_2_PtCl_6_ · 6H_2_O, 0.01% of lead acetate Pb(COOCH_3_)_2_ · 3H_2_O and 0.0025% of HCl [21]. An LCR meter (4263A, Hewlett Packard) was used for deposition at 800 mV and 100 Hz.

### 2.4 Electrochemistry

Electrochemical impedance spectroscopy (EIS) was performed with a potentiostat (CompactStat, Ivium Technologies). The arrays were soaked in phosphate-buffered saline (pH 7.4) at room temperature with a Pt counter wire and an Ag/AgCl reference wire. The potential was set at 50 mV, and the impedance magnitude and phase were measured between 1 Hz and 1 MHz. Cyclic voltammetry (CV) was performed with the same setup and three-electrodes configuration used for EIS. The applied voltage was scanned between −0.6 and 0.8 V at 50 mV s^-1^. CV was repeated for six cycles. The first cycle was discarded, and the remaining five were averaged.

### 2.5 Animal handling and surgery

Animal experiments were performed according to the authorisation GE519 approved by the Département de l’Emploi, des Affaires Sociales et de la Santé, Direction Générale de la Santé of the Republique et Canton de Genève (Switzerland), as previously described [17]. Two female Chinchilla Bastard rabbits (>16 weeks, >2.5 kg) were premedicated 30 min before the transfer to the surgical room with an intramuscular injection of xylazine (3 mg kg^-1^; Rompun®20 mg ml^-1^, 0.15 ml kg^-1^), ketamine (25 mg kg^-1^; Ketanarkon^®^ 100 mg ml^-1^, 0.25 ml kg^-1^) and buprenorphine (0.03 mg kg^-1^; Temgesic® 0.3 mg ml^-1^, 0.1 ml kg^-1^). Rabbits were placed on a heating pad at 35 °C. A 22G catheter was placed in the ear marginal vein 15 min after premedication. Local anaesthesia was provided to the throat (Xylocaine 10%, spray push). The rabbit was intubated with an endotracheal tube with balloon (3.5 mm) and ventilated (7 ml kg^-1^, rate: 40 min^-1^, positive end-expiratory pressure: 3 cm of H_2_O). Eye gel was placed on the eye to protect it from drying. Anaesthesia and analgesia were provided intravenously with propofol (10 mg kg^-1^ h^-1^; 20 mg ml^-1^, 0.5 ml kg^-1^ h^-1^) and fentanyl (0.005 mg kg^-1^ h^-1^; 0.05 mg ml^-1^, 0.1 ml kg^-1^ h^-1^). Body temperature, heart rate, blood pressure, and oxygen saturation were monitored continuously during the procedure. Saline was administered intravenously to prevent dehydration. The rabbit head was shaved and secured to a stereotactic frame (David Kopf Instruments). Lidocaine (6 mg kg^-1^; Lidocaine 20 mg ml^-1^, 0.3 ml kg^-1^) was injected subcutaneously at the surgical sites. After 5 min, the skin was opened and pulled aside to clean the skull with cotton swabs. A craniotomy was made to expose the visual cortex. The dura was removed, and an electrocorticography (ECoG) array was placed over V1 (E32-1000-30-200; NeuroNexus). The ECoG array was composed of 32 (4 × 8) Pt electrodes with a 200-μm diameter and a 1-mm pitch. A temporal craniotomy was made to access the optic nerve. The Flat-OpticSELINE was inserted in the intracranial portion of the nerve by piercing the nerve with a needle (nylon black DLZ 4,8-150 10/0, FSSB) and guiding the Flat-OpticSELINE transversely into the nerve. Rabbits were euthanised at the end of the procedure with an intravenous injection of pentobarbital (120 mg kg^-1^).

### 2.6 Electrophysiological recording and stimulation

The ECoG array was connected to the amplifier using a 32-channel analogue head stage for cortical recordings (PZ5; Tucker-Davis Technologies). The Flat-OpticSELINE was attached to a current stimulator (IZ2MH; Tucker-Davis Technologies). Optic nerve stimulation was performed with asymmetric biphasic cathodic-first current pulses (cathodic phase 150 μs; anodic phase 750 μs at one-fifth of the cathodic amplitude; interphase gap 20 μs) at various cathodic current amplitudes (10, 25, 50, 75, 100, 150, 200, 250, 500, 750, 1000, 1500 and 2000 μA). Each stimulus was composed of a single current pulse or a pulse train of 2, 3 or 4 pulses delivered at 1 kHz. Within the train, pulses had the same current amplitude. 52 conditions were generated by combining the current amplitudes (13) and the number of pulses in the train (1, 2, 3 or 4) for each of the 8 stimulating electrodes. A total of 416 stimuli were delivered in a randomised manner. The stimulation protocol was repeated 30 times (Rabbit 1) and 15 times (Rabbit 2) for both monopolar and 3D multilayer CB configurations.

### 2.7 Signal processing

Electrically evoked cortical potentials (EEPs) were filtered from 3 to 100 Hz and sampled at 24 kHz. Then, epochs synchronous to the stimulus onset (from −200 to +300 ms) were extracted from the data stream. Before data analysis in MATLAB, EEPs were visually inspected and manually rejected when presenting noisy signals. Afterwards, epochs were down-sampled to 2.4 kHz. An unsupervised approach based on Gaussian Mixture Model (GMM) clustering was used to classify electrodes exhibiting significant EEPs. For each rabbit, 25-ms long epochs after the stimulus onset were concatenated to perform dimensionality reduction with principal components analysis (PCA). The first two PCs were chosen to run the GMM clustering algorithm. The Davies-Bouldin index was used to select the optimal number of clusters. Templates for each cluster were plotted, and those presenting a flat signal were classified as electrodes not exhibiting a significant EEP (non-responsive electrodes). Only recording electrodes showing a significant EEP (responsive electrodes) were considered for further analysis. EEP peak-to-peak amplitudes were computed as the difference between P1 and N1 peaks. A normalised weighted distance was calculated using equation (1) to quantify how significantly responses spread across the visual cortex when changing the stimulus parameters [22]:

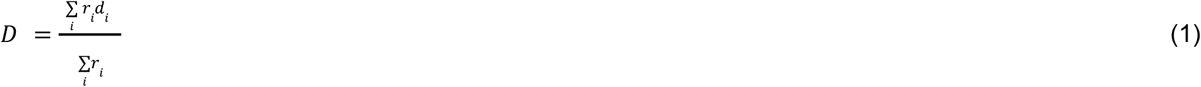

Where *r_i_* is the EEP peak-to-peak amplitude in responsive electrode *i*, and *d_i_* is the cortical distance in mm from the responsive electrode 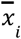 to the weighted mean centre 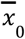 among the responsive electrodes, as in equation (2). *r_i_* is zero for the non-responsive electrodes.

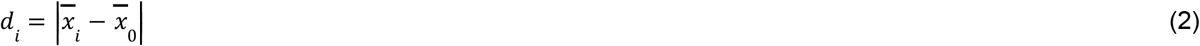

### 2.8 Statistical analysis and graphical representation

Statistical analysis and graphical representation were performed with Prism 8 (Graph Pad). The D’Agostino & Pearson omnibus normality test was performed to justify the choice of the statistical test. In plots, p-values were reported as: * p < 0.05, ** p < 0.01, *** p < 0.001, and **** p < 0.0001.

## 3. RESULTS

### 3.1 Hybrid FEA-NEURON modelling

Monopolar configuration is predominantly used in neural stimulation (**Figure 1b,c**). In this case, each electrode has a common and distant return electrode placed several millimetres away. Upon current injection, the electric field spreads from the stimulating electrodes to the common and distant return ground, resulting in an unconfined electrical activation of the neural tissue. We modelled by FEA a 5 x 5 microelectrode array (80-μm diameter and 100-μm pitch) in monopolar configuration. The simulation confirmed that the electric field spread widely (**Figure 1d,e**). Therefore, we designed a CB configuration with a 3D feature to achieve focal stimulation (**Figure 1f,g**). The central and the concentric return electrodes are placed in a multilayer configuration on two layers separated by 6 μm [15]. Moreover, they are independently controlled and referenced to a distal ground. The central electrode is the stimulating electrode (cathodic first), and the return injects a current of opposite polarity (anodic first). Therefore, the electric field is localised from the central electrode to the concentric electrode working as a local independent return. The return coefficient α (from 0.5 to 1) defines the proportion between the current injected into the central electrode, and the current returned to the concentric return. For α = 0.5, the CB stimulation is balanced, and the total current is returned to the concentric electrode. For α > 0.5, only a fraction of the current is returned to the concentric electrode, while the remaining current is returned to the distant ground (I_g_). Finally, for α = 1, the system works in the monopolar configuration, and the total current is returned to the distant ground (I_g_ = I). FEA simulations confirmed that the local return electrode localised the electric field (**Figure 1h,i**).

To further analyse the performance of CB electrodes and ensure that neighbouring electrodes do not have crosstalk, we repeated the simulation activating different electrodes of the array: central electrode only (**Figure 2a**, left), all electrodes (**Figure 2a**, middle) or all electrodes but with the central electrode off (**Figure 2a**, right). The electric potential profiles were obtained along the midline (dashed black line in **Figure 2a**) in planes at increasing distances from the surface of the array (sketch in **Figure 2a**, middle). When only one electrode is active, the electric potential profile obtained by FEA simulations has a Gaussian shape for both the monopolar configuration (**Figure 2b**, left) and the 3D multilayer CB configuration (**Figure 2c**, left). However, for the monopolar configuration, the lateral spreading of the electric potential increases as the distance from the electrode increase (**Figure 2b**, left), while it remains sharp for the 3D multilayer CB configuration (**Figure 2c**, left) even on a plane located 100 μm away from the electrode. When all the 25 electrodes are activated, the electric potential profile obtained for the monopolar configuration (**Figure 2b**, middle) is widened and increased in amplitude due to potential summation. The contribution of each electrode is barely distinguishable only at a distance of 10 μm from the electrode surface. Further away, the electric potential fuse. When switching off the central electrode, the electric potential in monopolar configuration remains blurred due to the electrode cross-talk (**Figure 2b**, right). The contrast between the central off electrode and the surrounding active electrodes may not be enough for discrimination. Thus, the electrodes in monopolar configuration cannot be considered electrically independent unless for a much lower density. Conversely, the 3D multilayer CB configuration allows sharp discrimination of the electric potential induced by each electrode at every distance (**Figure 2c**, middle). Also, we observed a maximal contrast in the electric potential drop when the central electrode is switched off (**Figure 2c**, right). In this condition, the electrodes are electrically independent, even for high electrode density.

**Figure 2.**
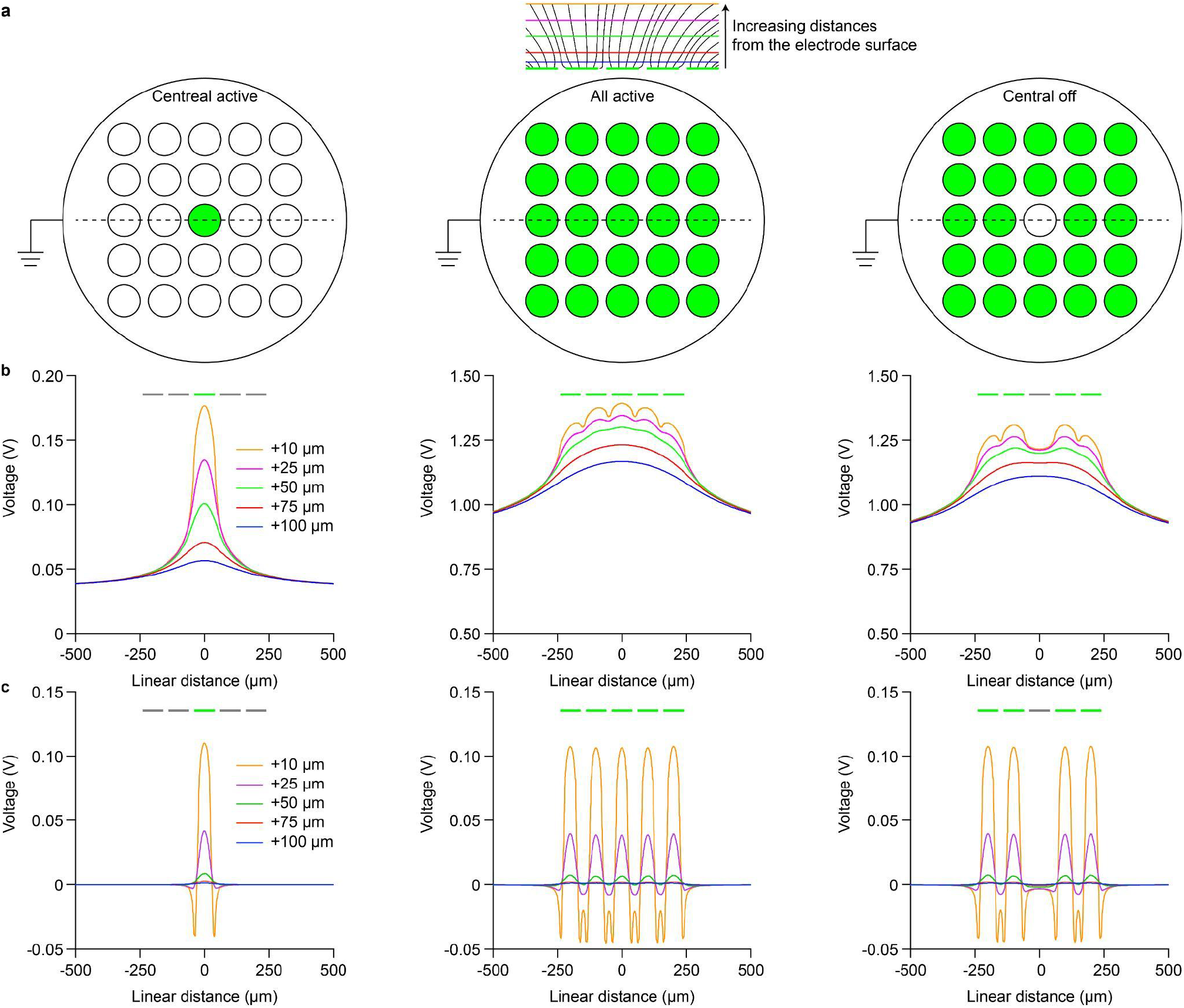
(**a**) Sketch of FEA simulations performed for three patterns: central electrode on (left), all electrodes on (middle) and all electrodes on but central electrode off (right). The electric potentials were measured along the dashed line and at various distances from the electrode surface: +10, +25, +50, +75, and +100 μm. (**b**) The plot of the electric potential obtained with the monopolar configuration and a current injected of I = −100 μA for the three patterns in (**a**). (**c**) The plot of the electric potential obtained with the 3D multilayer CB configuration (α = 0.5) and a current injected of −50 μA in the central electrode and +50 μA in the return electrode for the three patterns in (**a**). In all panels, the green/grey bars show the position of the active (green) and inactive (grey) electrodes.

FEA simulations showed that the 3D multilayer CB configuration focuses the electric potential laterally and vertically. The electric potential decreases substantially with the distance from the electrode surface. This decrease is due to the lateral shunting effect of the concentric return electrode. Thus, the stimulation is localised vertically to a few tens of micrometres from the electrode surface, which is not the case for the monopolar configuration. This feature is useful when a small area around the electrode has to be stimulated, such as optic nerve stimulation for artificial vision. However, measuring the recruitment of nerve fibres in-vivo is a challenging task. Therefore, we implemented a hybrid FEA-NEURON model. First, the electric field generated by the electrical stimulation was computed using FEA simulation for the monopolar and the 3D multilayer CB configuration (α = 0.5). For a better comparison, the dimension of the central cathode was 40 μm in both configurations. Thus, the monopolar configuration is equivalent to the 3D multilayer CB configuration with α = 1. Then, the activation probability of the fibres upon electrical stimulation for one electrode located in the centre of the nerve was obtained in NEURON. The model showed that the nerve area activated by one electrode in the 3D multilayer CB configuration (α = 0.5) is more confined than the monopolar configuration (**Figure 3a**). Besides the size of the activated area, also the shape is different between the two conditions. The activated area is circular around the central electrode in the monopolar configuration. For low current amplitudes, the shape is not symmetrical to the horizontal plane because the electrode is exposed towards the upper part of the nerve, and the PI substrate reduces the stimulation in the lower part of the nerve. At higher current amplitudes, the activation area becomes more symmetric because of the large spreading of the electric field. In the 3D multilayer CB configuration, the localisation of the electric potential due to the concentric return electrode blocks the activation of the nerve below the plane of the electrode array. This result is crucial to enhance selectivity in TIME arrays. Usually, TIME arrays have electrodes on both sides [11,16], and 3D multilayer CB electrodes can provide lateral selectivity of the stimulated area even at high current amplitudes. Interestingly, the activated area has a ‘three-leaf clover’ shape for 3D multilayer CB electrodes. This shape is caused by the anodic activation from the concentric return electrode. The quantification of the total number of activated fibres revealed that a 3D multilayer CB electrode activated about 40 times fewer fibres than a monopolar electrode (**Figure 3b,c**). Also, for 3D multilayer CB electrodes, we found that the return coefficient α linearly modulates the number of activated fibres (**Figure 3d**).

**Figure 3.**
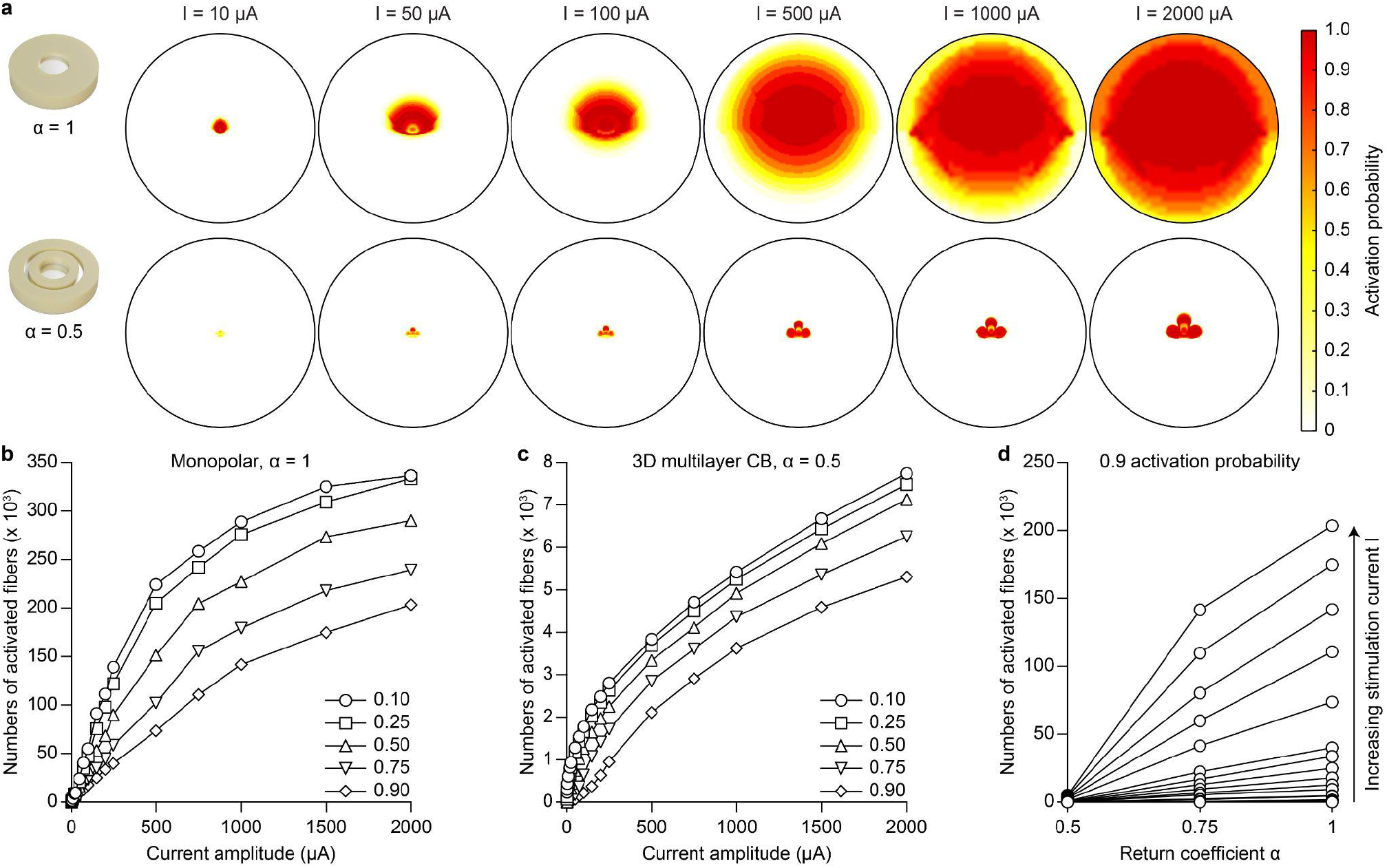
(**a**) Probability activation maps in the optic nerve upon current pulses at increasing current amplitudes from a monopolar electrode (α = 1, top) and a 3D multilayer CB electrode (α = 0.5, bottom). The black circles correspond to the optic nerve (diameter of 1500 μm). (**b,c**) The number of fibres activated by a single pulse as a function of the current amplitude from a monopolar electrode (**b**) and a 3D multilayer CB electrode (**c**). The quantification was made for 5 activation probabilities (higher than 0.1, 0.25, 0.5, 0.75 and 0.9). (**d**) The number of fibres activated with a probability higher than 0.9 by a single current pulse as a function of the return coefficient α for increasing current amplitudes. Each set of connected circles refers to one current amplitude. Simulated current amplitudes are 2, 3, 5, 10, 20, 25, 50, 75, 100, 150, 200, 250, 500, 750, 1000, 1500 and 2000 μA.

To further assess the higher selectivity in 3D multilayer CB electrodes, we compared two current amplitudes having a similar total activation probability across the entire nerve section (4.8% percentage change between the two configurations), which is the sum of all the values in the activation probability matrix sampled with a resolution of 1 x 1 μm^2^. The amplitude is 20 μA for the monopolar configuration and 2000 μA for the 3D multilayer CB (**Figure 4a**). It must also be clarified that a similar total activation probability corresponds to a similar number of activated fibres. Then, we computed the cumulative distribution of all the activation probabilities within the two maps (**Figure 4b**). The 3D multilayer CB configuration (red line) contains a higher proportion of high activation probabilities than the monopolar configuration (black line), visible from the left shift of the curve and the steeper increase. For instance, in the 3D multilayer CB configuration, 60% of the area is activated at a probability higher than 0.9 (dashed grey line), while only 40% in the monopolar configuration. This result shows that when the same averaged number of fibres are activated (similar total activation probability across the entire nerve section), the 3D multilayer CB configuration activates fibres (both small and large) in a smaller area close to the electrode. On the contrary, the monopolar configuration activates a more diffuse area. Indeed, despite the total activation probability across the entire nerve section being similar (4.8% percentage change), the total activated area (black contour in the maps) is larger by a 30.4% percentage increase in the monopolar configuration. The peripheral area with low activation probabilities visible in the monopolar configuration is associated with the activation of larger fibres away from the electrode since they have a lower activation threshold compared to smaller fibres.

**Figure 4.**
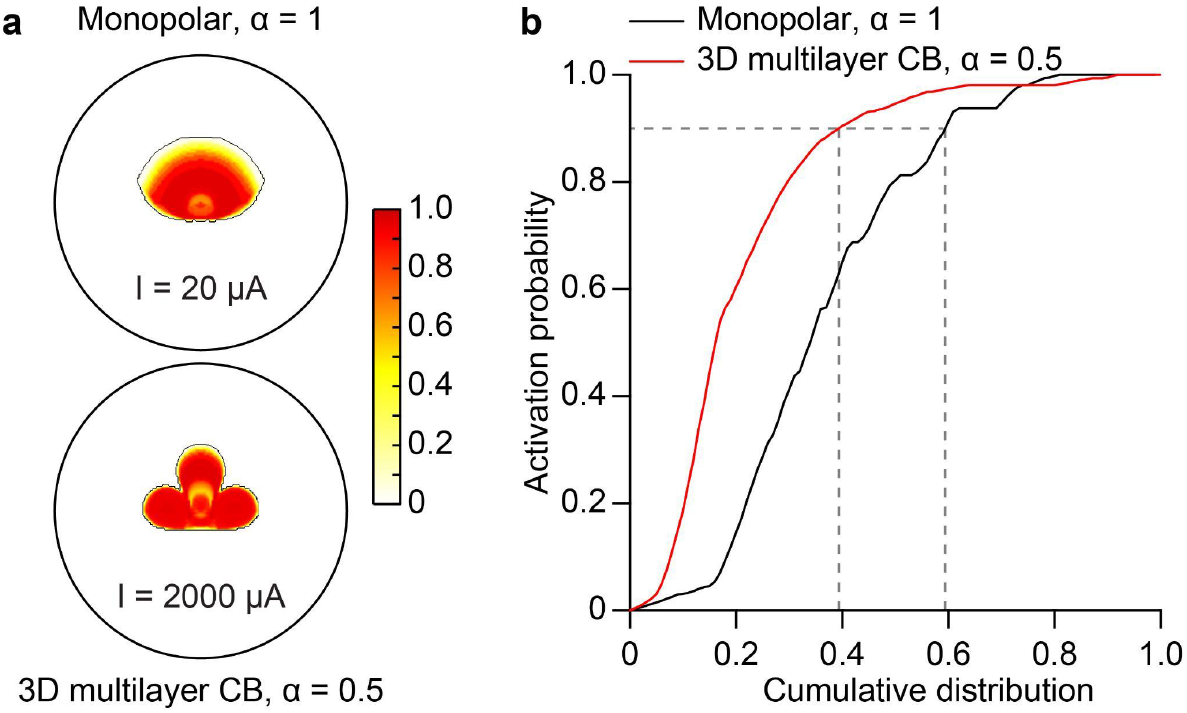
(**a**) Magnified view of two activation maps, corresponding to 20 μA for the monopolar electrode (α = 1, top) and 2000 μA for the 3D multilayer CB electrode (α = 0.5, bottom). The black contour delimits the total activated area (non-zero activation probability). The black circles correspond to a portion of the nerve with a diameter of 750 μm. The colour scale is the activation probability. (**b**) Plot of the activation probability as a function of the cumulative distribution of the non-zero activation probabilities within the activation matrix for the monopolar (α = 1, black) and the 3D multilayer CB (α = 0.5, red) configurations. Dashed grey lines highlighted the cumulative distribution at an activation probability of 0.9.

### 3.2 Flat-OpticSELINE microfabrication and characterisation

We fabricated a Flat-OpticSELINE electrode array with 8 3D multilayer CB electrodes having a 40-μm diameter and a 160-μm pitch (**Figure 5a**). Electrodes are placed in two layers to reduce the space occupied by the feedlines: the central electrodes are in the bottom layer (**Figure 5a**, grey), and the concentric return electrodes are in the top layer (**Figure 5a**, black). Feedlines from the same electrode pair are overlaid.

**Figure 5.**
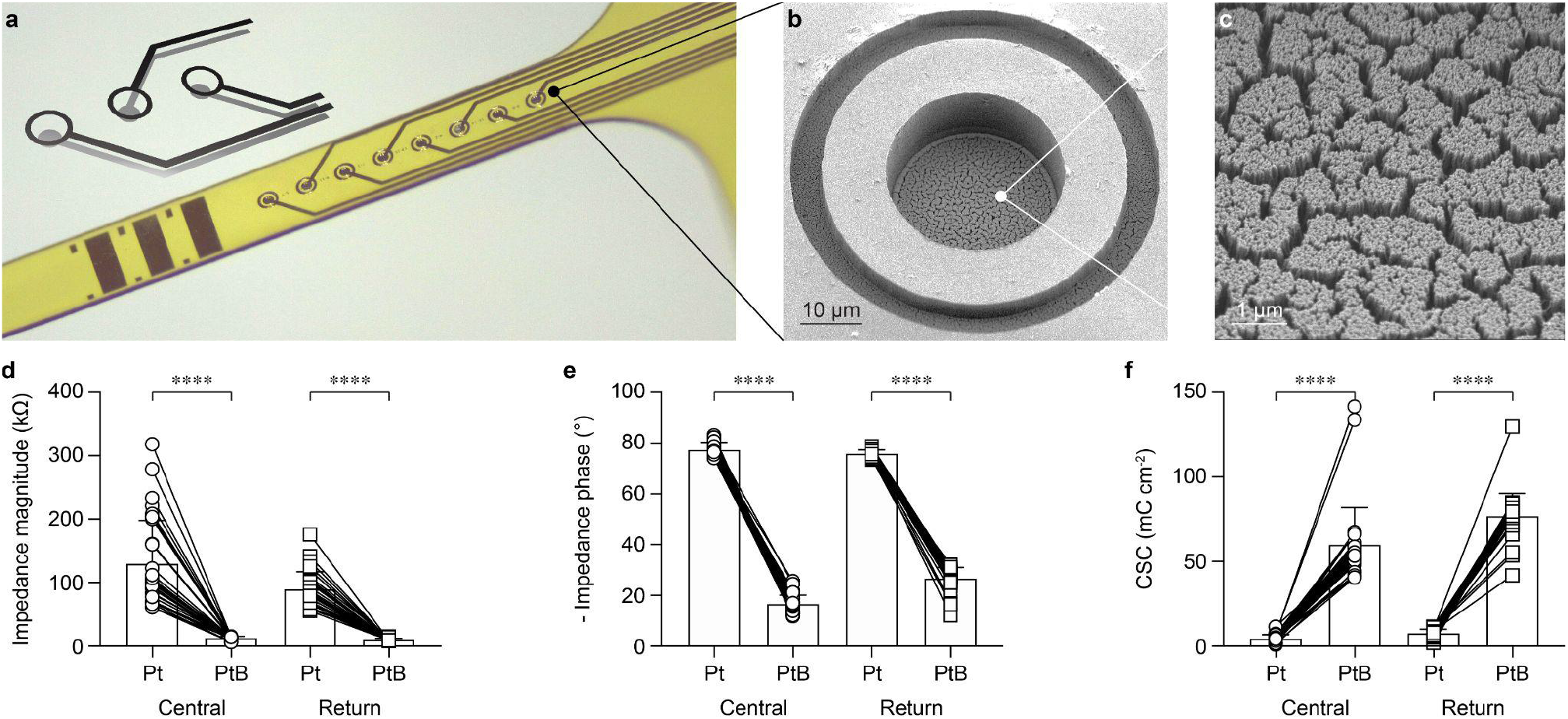
(**a**) Picture of a Flat-OpticSELINE with 8 3D multilayer CB electrodes. The sketch shows the electrode positions in the two planes. Central electrodes are in grey, and concentric return electrodes are in black. (**b**) Scanning electron microscope image of one 3D multilayer CB electrode. (**c**) Magnified scanning electron microscope image of the PtB coating. (**d**) Quantification of the impedance magnitude at 1 kHz before (circles) and after (squares) coating with PtB for both the central and return electrodes. (**e**) Quantification of the impedance phase at 1 kHz before (circles) and after (squares) coating with PtB for the central and return electrodes. (**f**) Quantification of the total charge storage capacity before (circles) and after (squares) coating with PtB for the central and return electrodes. In panels **d**-**f**, bars show the mean ± s.d.

Electrodes were coated with PtB (**Figure 5b,c**). EIS and CV showed that the impedance magnitude and phase at 1 kHz decreased (p < 0.0001 for both, two-tailed Wilcoxon matched-pairs signed rank test) and the charge storage capacity (CSC) increased (p < 0.0001, two-tailed Wilcoxon matched-pairs signed rank test) for both central and return electrodes after coating with PtB (**Figure 5d-f**).

### 3.3 Electrophysiological validation in-vivo

The Flat-OpticSELINE with 3D multilayer CB electrodes was transversally inserted in the optic nerve, and EEPs were recorded upon electrical stimulation with biphasic charge-balanced asymmetric current pulses of different current amplitudes ranging from 10 to 2000 μA (**Figure 6a**). EEPs were detected using an ECoG electrode array placed over V1 contralateral to the implanted optic nerve. In each experiment, EEPs were recorded from both monopolar (α = 1) and the 3D multilayer CB (α = 0.5) configurations (**Figure 6b,c**). After dimensionality reduction through PCA, significant responses to electrical stimulation were classified using a GMM clustering algorithm (**Figure 6d**). For each cluster, we then plotted the template responses. Clusters that appeared near the origin of the axes in the PCs space contained those responses that exhibited no stimulus-related modulation and showed a flat template (**Figure 6d**, red). Therefore, the other clusters were considered to contain significant responses (**Figure 6d**, green and blue). Last, we applied a detection algorithm to determine N1 and P1 peaks and compute the EEP peak-to-peak amplitudes.

**Figure 6.**
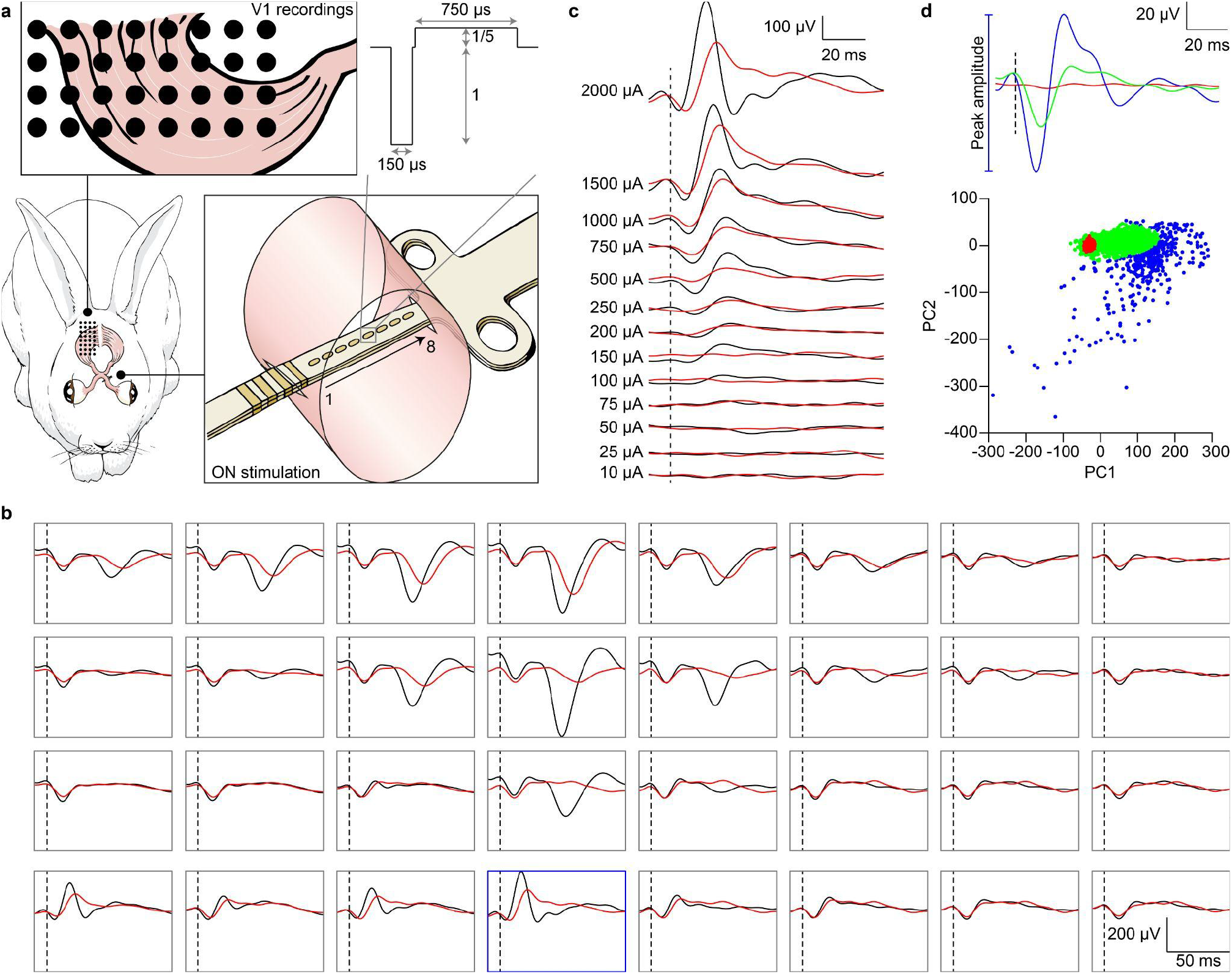
(**a**) Sketch of the experiment. (**b**) Representative examples of EEPs recorded upon monopolar (α = 1, black) and the 3D multilayer CB (α = 0.5, red) stimulations from the same electrode at the maximum current amplitude (2000 μA). The 4 × 8 boxes correspond to the arrangement of the 32 electrodes in the ECoG array. The dashed black lines show the onset of the electric stimulus. The blue box identifies the recording channel highlighted in panel **c**. (**c**) Representative example of EEPs recorded by one electrode (blu box in **b**) upon monopolar (α = 1, black) and the 3D multilayer CB (α = 0.5, red) stimulations at increasing current amplitudes. The dashed black line shows the onset of the electric stimulus. (**d**) Bottom: PC1-PC2 space after GMM Clustering (k = 3). Top: template responses from each cluster.

The mean EEP peak-to-peak amplitudes for all the recording and stimulating electrodes in both conditions, monopolar (α = 1) and 3D multilayer CB (α = 0.5), showed a monotonic growth as a function of the current amplitude (**Figure 7a**). It was also apparent that EEP peak-to-peak amplitudes became higher than zero, starting from a current amplitude of 150 μA for the monopolar configuration and 250 μA for the 3D multilayer CB. A linear regression fitting showed that the slopes of the two lines are significantly different (p < 0.0001), confirming the prediction of the hybrid FEA-NEURON model. Next, we assessed at which current amplitude each stimulating electrode elicited a significantly different averaged EEP peak-to-peak amplitude between the monopolar (α = 1) and the 3D multilayer CB (α = 0.5) configurations using the Kolmogorov-Smirnov test (**Figure 7b**). From 150 μA, an increasing number of stimulating electrodes appeared to elicit a significantly different EEP. While from 1500 μA, all the stimulating electrodes (8 per rabbit) showed a significant difference between monopolar (α = 1) and the 3D multilayer CB (α = 0.5) configurations.

**Figure 7.**
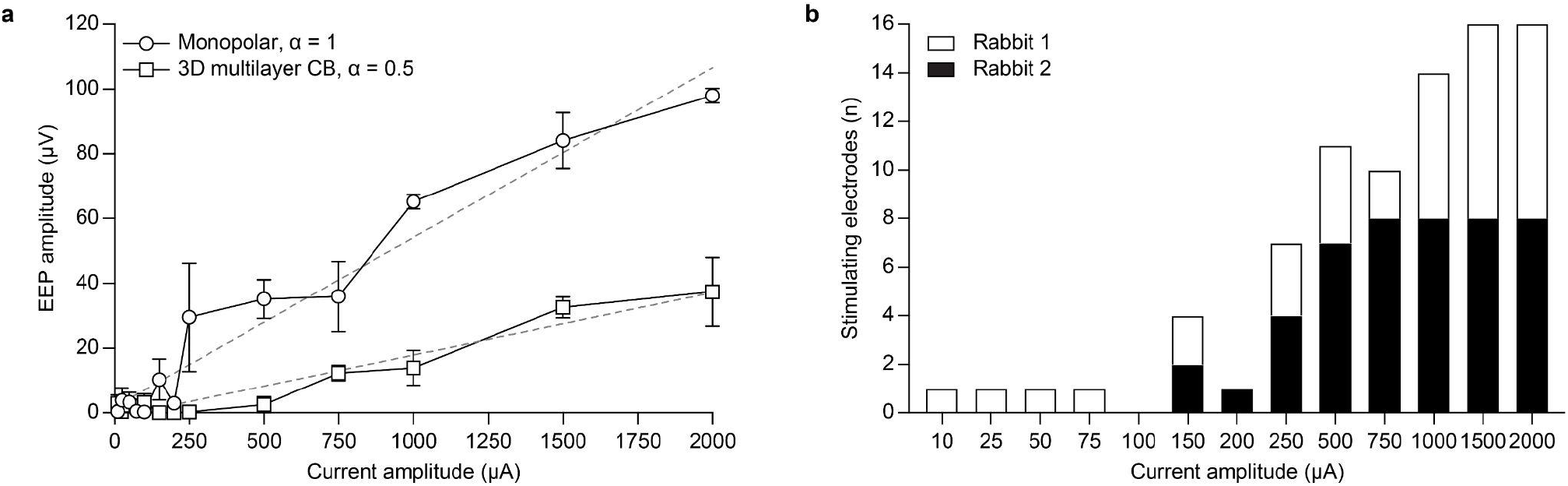
(**a**) Quantification of the mean (± s.e.m., N = 2 rabbits) EEP peak-to-peak amplitudes for all the recording and stimulating electrodes in monopolar (α = 1, circles) and the 3D multilayer CB (α = 0.5, squares) configurations. The dashed grey lines are linear regressions. For monopolar configuration: slope = 0.0524, R squared = 0.9140. For 3D multilayer CB configuration: slope = 0.0192, R squared = 0.8488. Both slopes are significantly non-zero (p < 0.0001). (**b**) Quantification of the number of stimulating electrodes that present a significant difference (p < 0.05, Kolmogorov-Smirnov test) in the averaged EEP peak-to-peak amplitude at different current amplitudes between monopolar (α = 1) and the 3D multilayer CB (α = 0.5) configurations.

The previous analysis compared the averaged EEP peak-to-peak amplitudes in both monopolar (α = 1) and 3D multilayer CB (α = 0.5) configurations, but it did not consider the spatial extent of the evoked response. Qualitatively, the evoked cortical response spread across more recording channels for increasing current amplitudes. However, it remained more localised when the 3D multilayer CB configuration was used instead of the monopolar configuration (**Figure 8a**). To quantify this difference, we computed two indexes: (1) the number of responsive electrodes (**Figure 8b,d**) and (2) the normalised weighted distance (**Figure 8c,e**). Results showed that fewer recording electrodes had a significant response in the 3D multilayer CB configuration compared to the monopolar configuration (**Figure 8b,d**). Polling the data of two rabbits together, one curve cannot fit both sets (p < 0.0001, F = 13.86, DFn = 4, DFd = 44) indicating that the two curves are significantly different (**Figure 8d**). Similarly, the normalised weighted distance was smaller, at every current amplitude, for the 3D multilayer CB configuration (**Figure 8c,e**). Polling the data of two rabbits together, one curve cannot fit both sets (p < 0.0001, F = 11.80, DFn = 4, DFd = 44) indicating that the two curves are significantly different (**Figure 8e**).

**Figure 8.**
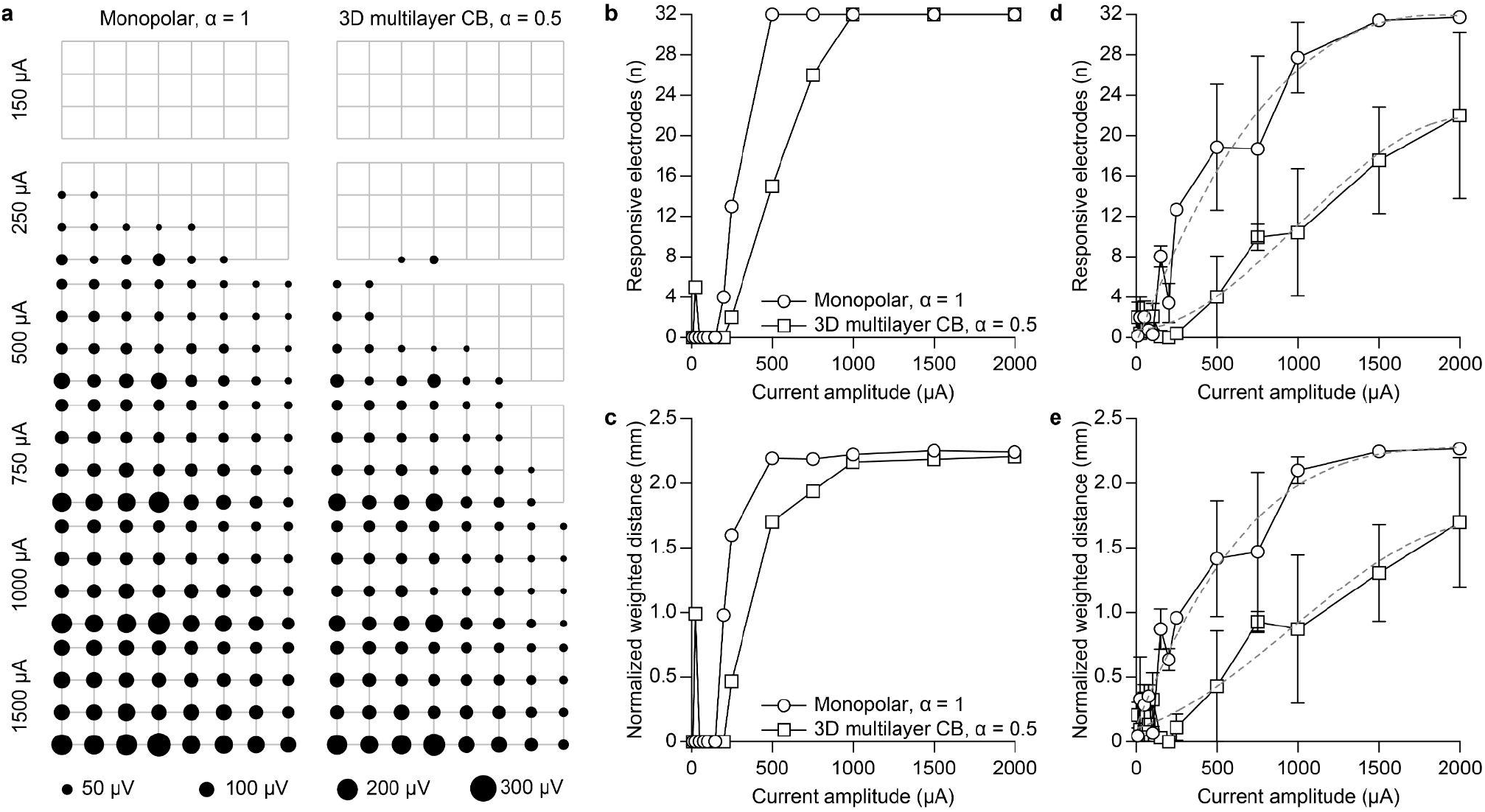
(**a**) Activation maps for one stimulating electrode in monopolar (α = 1) and the 3D multilayer CB (α = 0.5) configurations. The size of the circles corresponds to the EEP peak-to-peak amplitudes. Only responsive electrodes are shown. The grey grid highlights the electrode positions. (**b**) Quantification of the number of responsive electrodes as a function of the current amplitude in monopolar (α = 1, circles) and the 3D multilayer CB (α = 0.5, squares) configurations. Representative example for one stimulating electrode in one rabbit. (**c**) Quantification of the normalised weighted distance as a function of the current amplitude in monopolar (α = 1, circles) and the 3D multilayer CB (α = 0.5, squares) configurations. Representative example for one stimulating electrode in one rabbit. (**d**) Quantification (mean ± s.e.m) of the number of responsive electrodes as a function of the current amplitude in monopolar (α = 1, circles) and the 3D multilayer CB (α = 0.5, squares) configurations across all the stimulation electrodes (n = 8 electrodes) and animals (N = 2 rabbits). The dashed grey lines are the third-order polynomial (cubic) regressions. For monopolar configuration: R squared = 0.8873. For 3D multilayer CB configuration: R squared = 0.7785. (**e**) Quantification (mean ± s.e.m) of the normalized weighted distance as a function of the current amplitude in monopolar (α = 1, circles) and the 3D multilayer CB (α = 0.5, squares) configurations across all the stimulation electrodes (n = 8 electrodes) and animals (N = 2 rabbits). The dashed grey lines are the third-order polynomial (cubic) regressions. For monopolar configuration: R squared = 0.8725. For 3D multilayer CB configuration: R squared = 0.7306.

Last, we investigated the possibility of modulating the cortical responsivity by changing the return coefficient α or increasing the number of pulses in the train (**Figure 9**). When changing the return coefficient, the linear regression fitting showed that the slopes of the two lines (α = 0.5 vs. α = 0.75) were significantly different (p = 0.0394 for 1 pulse; p < 0.0001 for 2 pulses; p < 0.0001 for 3 pulses; p < 0.0001 for 4 pulses). This result confirms the prediction of the hybrid FEA-NEURON model (**Figure 3d**). Similarly, when increasing the number of pulses in the train, the linear regression fitting showed that the slopes of the four lines were significantly different (p = 0.0035 for α = 0.5; p < 0.0001 for α = 0.75). This result confirms the previous finding that the cortical activation can be modulated in amplitude by increasing the number of pulses instead of changing the current intensity [16]. More pulses in the train increased the EEP peak-to-peak amplitude by increasing the probability of fibre activation via a mechanism of temporal summation resulting from the repeated activation of the same few fibres and not by the enlargement of the activated area of the nerve.

**Figure 9.**
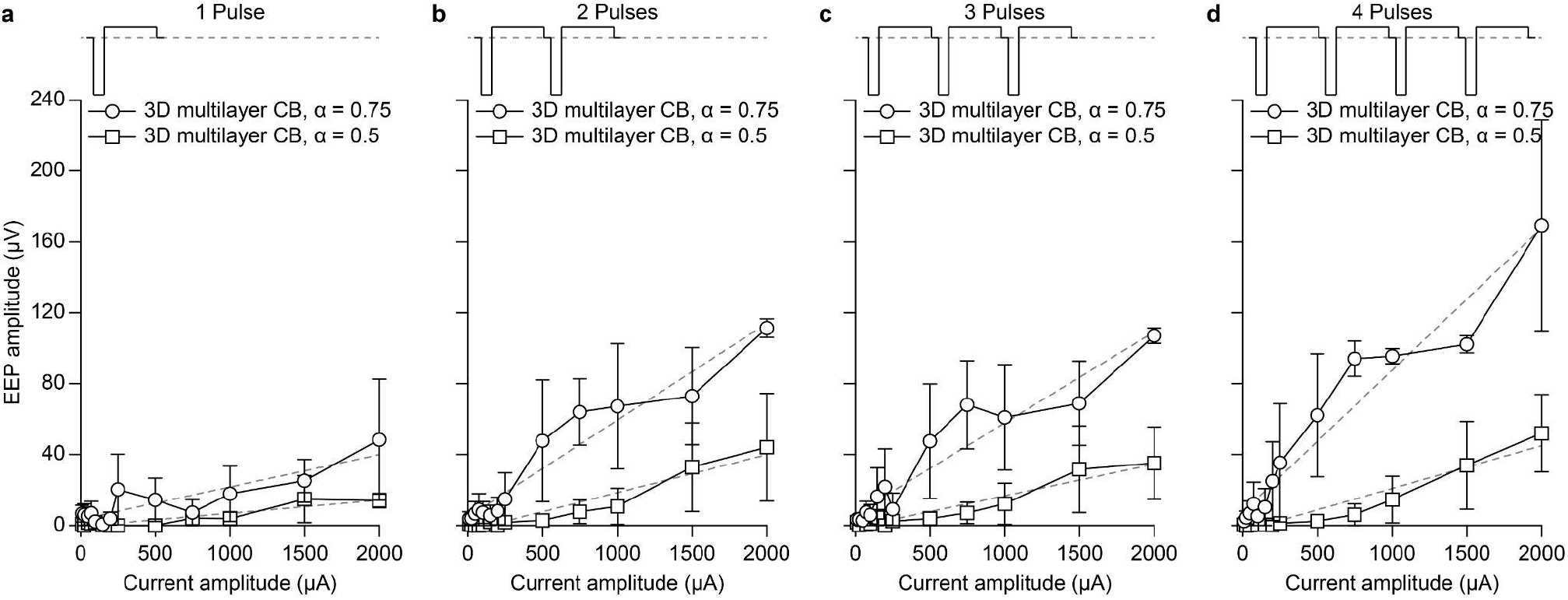
Quantification (mean ± s.e.m., N = 2 rabbits) of the EEP peak-to-peak amplitudes across all the recording and stimulating electrodes in the 3D multilayer CB configuration with return coefficient α = 0.75 (circles) and α = 0.5 (squares) for trains with 1 pulse (**a**), 2 pulses (**b**), 3 pulses (**c**) and 4 pulses (**d**) at 1 kHz pulse rate. The dashed grey lines are linear regressions. All slopes are significantly non-zero (p < 0.0001).

## 4. DISCUSSION

In this article, we proposed a 3D multilayer CB electrode array that enhances the selectivity of optic nerve stimulation. We first investigated the effectiveness of 3D multilayer CB configuration in reducing the number of fibres activated by electrical stimulation with a hybrid FEA-NEURON model. Then, we assessed in-vivo the spread of neural activation in the visual cortex compared to monopolar stimulation. These findings are consistent with previous evidence that local return stimulation can be used to restrict the electrical activation of neurons by shaping the electric field [23,24].

Other approaches have been proposed to localise electrical stimulation. A first approach, called quasi-monopolar (QMP), has a distant ground electrode and a plane of hexapolar return electrodes surrounding each stimulating electrode. Each electrode in the array is alternatively used as a stimulating electrode or as part of the hexapolar return. This approach has been implemented in a suprachoroidal prosthesis under development by Bionic Vision Technologies [13,25], but several disadvantages have been identified [26]. QMP requires complex current injection schemes at each of the hexapolar electrodes, and the electrodes in the external perimeter of the array cannot be used for QMP stimulation. Also, the QMP configuration is suitable for large surface electrode arrays (e.g. retinal prostheses or ECoG arrays) and not for intraneural electrode arrays. Last, complex stimulation patterns at high resolution cannot be easily obtained since, for every stimulating electrode, the six surrounding electrodes are used as a return. A second option is the ground plane configuration, where a highly conductive local ground plane provides focalisation of the currents [27,28]. Compared to the QMP configuration, its main advantage is that it requires fewer feedlines, which is necessary for arrays with many electrodes (e.g. > few hundreds). Besides this advantage, it has several limitations. First, it requires a highly conductive surface (> 5,000 S m^-2^). A metallic ground plane exposed to the extracellular medium is not advantageous because of the risk of delamination, it makes the device not transparent (if necessary), and it may crack due to the implant bending necessary in most surgical insertions. Last, a ground plane surrounding each electrode of the array limits the final electrode density.

A 3D multilayer CB electrode provides a stimulation configuration that strongly reduces crosstalk between electrodes. The multilayer process allows placing concentric electrodes on different layers and patterning the feedlines from the same electrode overlaid. Therefore, each electrode pair occupies the same space on the array as the monopolar configuration. However, because the two electrodes are independently controlled, the number of pads will be doubled compared to the monopolar configuration.

In the study, we compared monopolar and 3D multilayer CB configurations using a hybrid FEA-NEURON model of the optic nerve. The results showed a substantial reduction in fibres activated with the 3D multilayer CB electrode. However, measuring the fibre activation upon electrical stimulation in the optic nerve remains an open challenge due to the limited space available in the intracranial segment, where the array is currently implanted. In addition to the stimulating array, another electrode array should be inserted for recordings to validate simulations. Future works may address this challenge ex-vivo combining stimulation and recording on explanted optic nerves [29]. Here, we choose to measure the activation in the visual cortex. Thus, we could infer that a lower peak-to-peak amplitude of the EEPs with the 3D multilayer CB configuration is associated with fewer fibres activated in the optic nerve.

To select significant EEPs, we first used PCA as a feature extraction method. We selected the first two principal components representing more than 95% of the variance [30]. EEPs were clustered according to a GMM algorithm: an unsupervised method used to cluster unlabeled data with advantages over the more classic k-means. The most important advantage is that GMM accounts for the data variance and can handle oblong clusters. The dimensionality reduction and clustering of the EEPs were automatic and relatively assumption-free [31], enabling a straightforward classification of significant EEP without defining a priori punctual features, such as latency or amplitude. Indeed, we already showed that punctual features appeared to be not informative enough for prediction models [17]. Hence, this classification allowed us to compare how significant electrically evoked responses spread across the cortex with changing stimulus parameters and electrode configuration.

Consistent with a previous study on suprachoroidal stimulation [32], the 3D multilayer CB configuration activated smaller EEPs and a smaller area on the cortex than the monopolar electrodes. We then quantified the activated area by computing the response spread across the visual cortex. We showed it increased with increasing current amplitudes for both monopolar, as presented in [22], and 3D multilayer CB configurations. However, on average, the extent of this spatial spread was significantly lower for the 3D multilayer CB configuration. Moreover, we also find that the current threshold for cortical activation with the 3D multilayer CB configuration was higher than for monopolar. It is possible that the signal-to-noise ratio and the spatial resolution of the ECoG array used in this study were not sufficient to record EEP upon activation of a small number of fibres, such as with low current amplitude in 3D multilayer CB configuration. Therefore, future works should employ recording solutions with higher spatial resolution. High-density intracortical microelectrodes arrays (e.g. Utah array) placed in multiple locations of the visual cortex could help assess the performance of the 3D multilayer CB configuration at low currents. Alternatively, fast three-dimensional functional imaging tools [33,34] could also provide better spatial resolution in cortical recordings.

Last, we observed that the modulation of the return coefficient α could increase the number of fibres activated and the EEP peak-to-peak amplitude. It must be noted that we designed 3D multilayer CB electrodes where the area of the central and return electrodes are almost identical. Thus, further work could investigate electrodes having different surface areas to optimise the stimulation efficiency.

## 5. CONCLUSION

Spatial selectivity in neuromodulation refers to interacting with neurons in a predefined and limited volume of tissue [35]. This ability is key in avoiding side effects and improving outcomes for neurological treatments. Highly localised electrical stimulation is an attractive solution to reach spatial selectivity. Here, we proposed a feasibility study of a 3D multilayer CB configuration carried out in the optic nerve. However, this approach could be extended to other applications. For example, in retinal prostheses, this configuration could help to increase spatial resolution. In cortical stimulation with surface electrodes, activation of a confined area in the somatosensory cortex could improve sensory feedback giving higher spatial discrimination in the corresponding perceived body parts [36]. A higher spatial resolution would be equally important in cortical visual prostheses, like the Orion I device [37]. Moreover, stimulation selectivity applies to peripheral nerve stimulation as well. For instance, it could help limit the side effects of unspecific vagus nerve stimulation and improve the treatment of pathological conditions [38,39]. Lastly, the 3D multilayer CB configuration could better modulate bladder functions in spinal cord injured patients [40]. In any case, spatially precise electrical stimulation requires a microscopic understanding of the anatomical and functional organisation of the target tissue, which is often missing, like for the optic nerve. This knowledge might provide a further step toward personalised bioelectronic medicine.

## 6. ACKNOWLEDGEMENT

This work was supported by École Polytechnique Fédérale de Lausanne, Medtronic plc and the Swiss National Science Foundation (200021_182670).

E.B. fabricated the devices, performed experiments and data analysis, and wrote the manuscript. V.G performed simulations. M.J.I.A.L. designed the fabrication process. E.G.Z. performed animal care and anaesthesia. R.C.M. performed simulations. D.G. led the study and wrote the manuscript. All the authors read and accepted the manuscript.

The authors declare no competing interests.

